# Cephalopod Sex Determination and its Ancient Evolutionary Origin

**DOI:** 10.1101/2024.02.21.581452

**Authors:** Gabrielle C. Coffing, Silas Tittes, Scott T. Small, Jeremea O. Songco-Casey, Denise M. Piscopo, Judit R. Pungor, Adam C. Miller, Cristopher M. Niell, Andrew D. Kern

## Abstract

Octopuses, squids, and cuttlefishes – the coleoid cephalopods – are a remarkable branch in the tree of life whose members exhibit a repertoire of sophisticated behaviors (Hanlon and Messenger, 2018). As a clade, coleoids harbor an incredible variety of novel traits including the most complex nervous system among invertebrates, derived camera-type eyes, and rapid adaptive camouflage abilities (Young, 1971; Hanlon, 2007). The burst of evolutionary novelty that distinguishes cephalopods is even more striking in a phylogenetic context; cephalopods are a deeply diverged lineage that last share a common ancestor with other extant molluscs in the Cambrian period, roughly 550 million years ago (Ponder and Lindberg, 2008; Huang et al., 2022). With recent advances in genome sequencing technologies, we have the capability to explore the genomic foundations of cephalopod novelties. Here, using PacBio long-read sequencing of genomic DNA and IsoSeq full-length mRNA sequencing, we provide a novel chromosome-scale reference genome and annotation for a female California two-spot octopus (*O. bimaculoides*). Our assembly reveals that the female octopus has just one sex chromosome, consistent with a ZO karyotype, while the male has two (ZZ), providing the first evidence of genetic sex determination in cephalopods. We use our assembly and annotation in combination with existing genomic information from other cephalopods to create the first whole genome alignments from this group and demonstrate that the sex chromosome is of an ancient origin, before the radiation of extant cephalopods approximately 480 million years ago (Huang et al., 2022), and has been conserved to the present day in all cephalopod genomes available.

## A chromosome-level assembly reveals a hemizygous Z chromosome in female octopus

The California two-spot octopus was the first cephalopod genome to be sequenced in 2015 (Albertin et al., 2015) and subsequently placed into scaffolds in 2022 (Albertin et al., 2022b). Although these resources have been tremendously valuable for cephalopod research, the assembly still contains numerous gaps and many gene annotations remain fragmented due to the highly-repetitive nature of the genome. To sequence through the long, repetitive stretches of the *O. bimaculoides* genome, we re-sequenced a single female individual with PacBio’s long, high-fidelity (HiFi) sequencing and used chromosomal conformation capture (Hi-C) to place scaffolds into chromosomes. After scaffolding, the total genome assembly size is 2.3Gb with 30 chromosomal scaffolds representing a *N* = 30 karyotype, which matches the findings of cytogenetic studies in other octopus species (Wang and Zheng, 2018). Additionally, we used IsoSeq sequencing to aid with annotating full-length genes. A comparison of assembly statistics to existing *O. bimaculoides* assemblies is shown in Table S1 and genome annotation statistics are in Table S2. Generally, our new assembly is more complete than previous assemblies, with a contig N50 of 0.86 Mb and a scaffold N50 of 101.05 Mb. The new assembly reduced the number of scaffolds in the assembly from 145,326 (Albertin et al., 2022b) to 583, increasing the average scaffold length by roughly three orders of magnitude.

Strikingly, our Hi-C contact map showed evidence for one chromosome, chromosome 17, having reduced coverage in comparison to other chromosomes in our assembly from a female individual (Figs. 1A & S1). As the original reference assembly from (Albertin et al., 2022b) was from a male individual we were able to compare coverage in that assembly, which showed no difference in coverage between chromosome 17 and any other chromosome. Using short read Illumina data that we generated from unrelated female and male *O. bimaculoides* individuals (*N* = 2 of each sex; Table S3) we confirmed that females of this species are hemizygous for chromosome 17 whereas males are diploid, so hereafter we refer to chromosome 17 as Z (Figs. 1B & S2).

**Figure 1.**
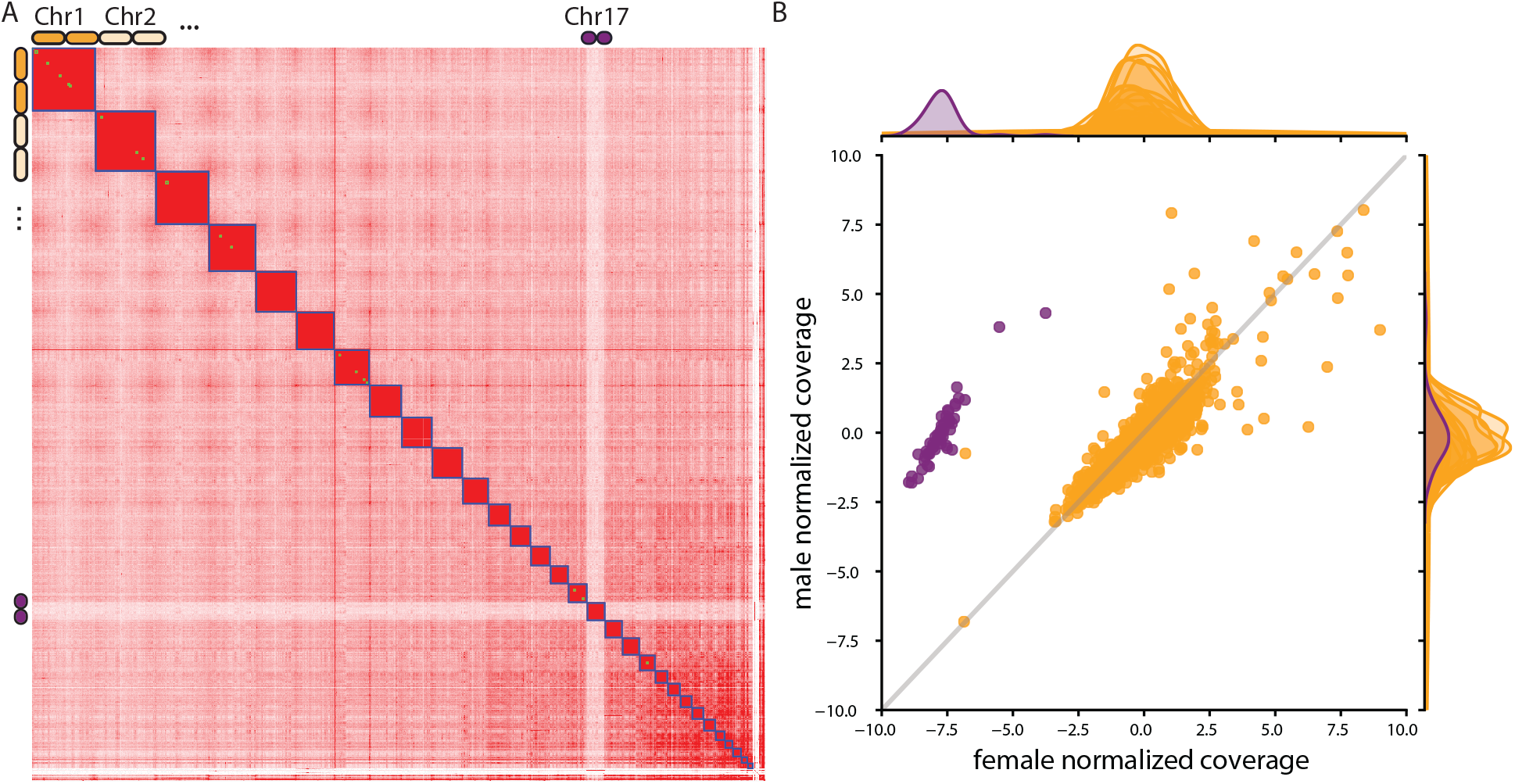
Sequencing data showing half coverage at chromosome 17 of *O. bimaculoides*. **A**. Hi-C contact map from our assembly of a female *O. bimaculoides*. Chromosome 17, our putative Z chromosome, is highlighted as a clear outlier in coverage. **B**. Normalized read depth of male and female whole-genome Illumina short read data. Purple points are from chromosome 17, whereas every other scaffold is shaded in orange. Scatterplot showing male vs. female normalized coverage, while the density estimates on the margins are from female (top) and male (side) separately.

Since the *O. bimaculoides* male genotype is clearly ZZ and the female is hemizygous at Z, we next turned our attention towards identification of a potential W chromosome limited to females. To do so we searched for scaffolds that were only present in the female-derived sequence libraries and absent from the male-derived libraries and assemblies. We found no such candidates, suggesting that females are ZO and males are ZZ. This perhaps indicates the evolutionary loss of the W chromosome after substantial degradation (Charlesworth et al., 2005; Charlesworth, 2013). It is possible that a small dot W exists that cannot be identified with our sequencing data. However, while ZZ/ZO sex determination systems are relatively rare among species found in the literature (Jonika et al., 2022), they have been described in several groups including Lepidopterans (Johnson and Lachance, 2012), plants (Charlesworth, 2013), amphibians (Perkins et al., 2019), and more (Bachtrog et al., 2014; Tree of Sex Consortium, 2014).

## Genomic comparisons among cephalopods show the Z chromosome is an outlier

To determine if the Z chromosome was unique to *O. bimaculoides* or more broadly distributed among cephalopods, we inferred a species tree of cephalopods using protein sequences with the STAG algorithm in OrthoFinder (Emms and Kelly, 2019). This alignment compared three other octopus genomes (*Hapalochlaena maculosa, Octopus minor*, and *Octopus sinensis*), two squid genomes (*Architeuthis dux*, and *Euprymna scolopes*), a cuttlefish genome (*Sepia esculenta*), and the outgroup to coleoid cephalopods, the chambered nautilus (*Nautilus pompilius*) (Fig. 2A). Of 237,704 proteins input to OrthoFinder, 166,258 (69.9%) were assigned to 18,791 orthogroups. Next, we created the first whole-genome, multiple alignment among existing cephalopod genomes using the Progressive Cactus / Comparative Genomics toolkit pipeline (Armstrong et al., 2020) (see Supplemental Methods for details). Our alignment enables a host of analyses, including studies of sequence divergence and synteny, and we were particularly interested to compare patterns of evolution on autosomes versus the Z chromosome.

**Figure 2.**
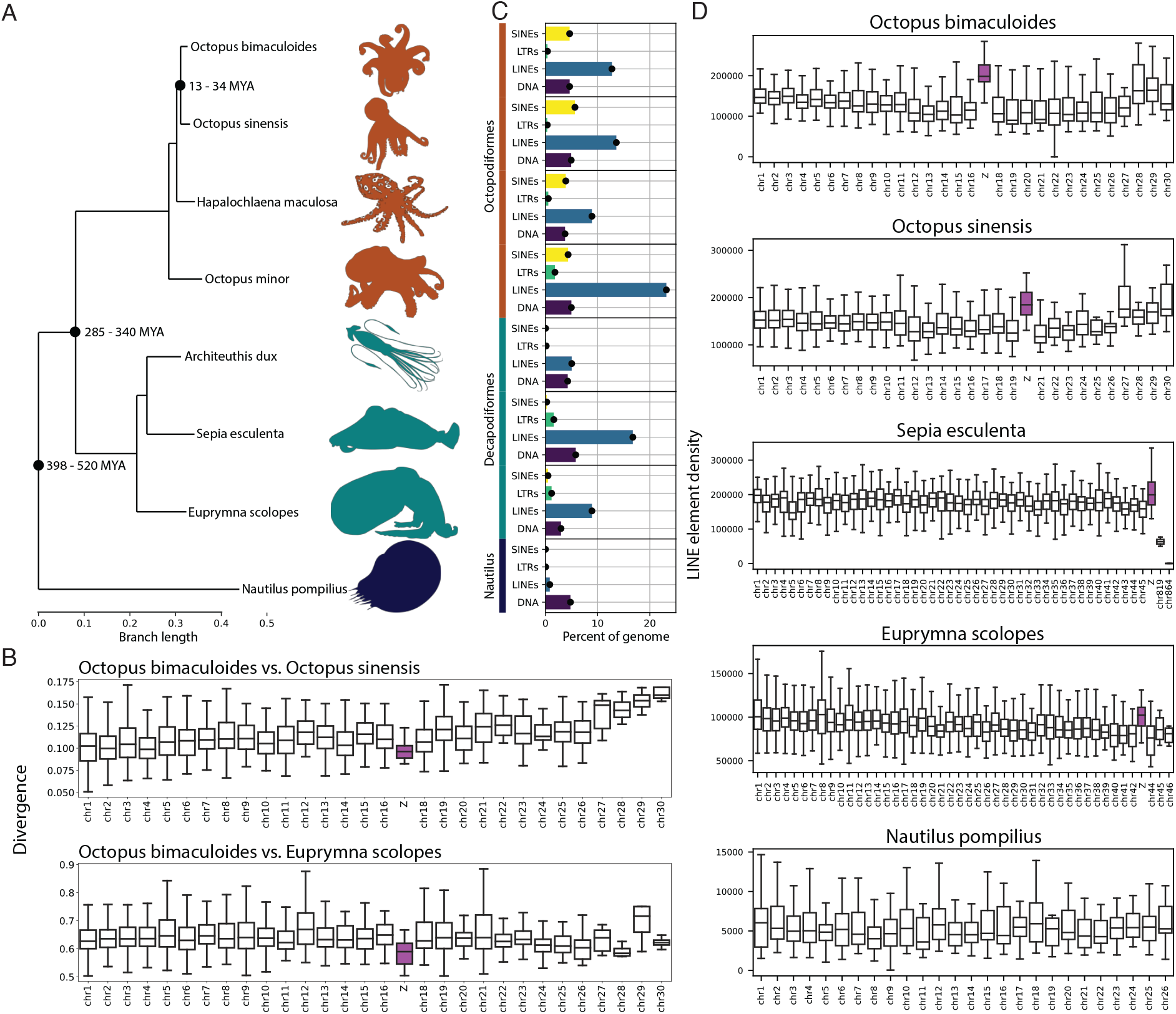
**A**. Cephalopod phylogeny used in this study. Phylogeny inferred using one-to-one protein orthologs identified by OrthoFinder Emms and Kelly (2019). Branch lengths in units of amino acid substitutions per site. Divergence dates are from (Huang et al., 2022). **B**. Windowed pairwise divergence calculated between *O. bimaculoides* and *O. sinesis* (top panel) and *O. bimaculoides* and *E. scolopes* (bottom panel). Divergence was calculated in 1Mb, non-overlapping windows using our multiple genome alignment with *O. bimaculoides* as the reference. The Zchromosome is highlighted in purple. **C**. Genomic repeats associated with each leaf taxon. Repeats show percent of genome composed of SINE elements (yellow), LTR elements (green), LINE elements (blue), and DNA transposons (purple) for each species. **D**. LINE content across genomes of five cephalopod species. The number of LINE base pairs were counted in 1Mb windows. In each of the coleoid species, we found a single chromosome with elevated LINE density. These putative Z chromosomes have been highlighted in purple. In *N. pompilius*, no chromosomes have outlying LINE densities.

We calculated divergence in 1 megabase windows between *O. bimaculoides* and *O. sinensis* using the phast package (GTR model) (Siepel et al., 2005), which revealed that the Z chromosome evolves significantly slower than autosomes (Mann–Whitney U test, p = 0.0011; Fig. 2B). The same pattern holds true for Z chromosome divergence between *O. bimaculoides* and the Hawaiian bobtail squid *Euprymna scolopes* (Mann–Whitney U test, p < 0.0001; Fig. 2B). We note that by using *O. bimaculoides* as a reference, the observed differences in divergence rates between Z and autosomes is likely conservative, as any loci which have moved moved between Z and autosomal chromosomes over evolutionary time should only obscure a difference, if present.

We also analyzed patterns of coding substitutions between *O. bimaculoides* and *O. sinensis*, restricting calculations to Z-linked genes in both species. While we found that the rate of nonsynonymous substitution was roughly equivalent between Z-linked and autosomal genes (Mann–Whitney U test, p = 0.38), Z-linked genes have decreased levels of synonymous substitution (mean Z-linked dS= 0.091, mean autosomal dS= 0.123; Mann–Whitney U test, *p*< 0.00001; Fig. S7). Reduced divergence on the Z mirrors what is observed in the primate X chromosome relative to autosomes (Hobolth et al., 2007; Scally et al., 2012), and may be caused by a variety of reasons including purging of recessive, mildly deleterious mutations from the hemizygous chromosome and sex-biased mutation rate differences between males and females. Using the ratio of the substitution rate at Z-linked versus autosomal loci allows us to estimate the degree of sex-biased mutation (Miyata et al., 1987), generally termed α. We find that for synonymous divergence α = 0.127 (95% C.I.= 0.022 − 0.268) meaning that for these positions we estimate the female mutation rate to be ∼ 7.8 times that of the male mutation rate. While synonymous substitutions are a decent proxy for neutral divergence, we can get a bulk estimate of α from the genome by simply considering divergence in large genomic windows. Using 1Mb windowed divergence yields an estimate of α = 0.539 (95% C.I.= 0.406*−*0.703) indicating that the female mutation rate is ∼ 1.82 times that of the male mutation rate genome-wide. This observation of female-biased mutation is notable and has been seen in fish lineages (Bergeron et al., 2023) but contrasts to the commonly-observed male bias in amniote lineages (Wilson Sayres and Makova, 2011; Bergeron et al., 2023). Although female-biased mutaiton rate is quite rare, *O. bimaculoides* females undergo synchronous ovulation (Ibáñez et al., 2021), a process that has been hypothesized to contribute to higher mutation rate in females (Bergeron et al., 2023).

## Repetitive element characteristics of the Z chromosome are unique

Our whole-genome alignment allowed us to examine repetitive element evolution in a phylogenetic framework. Figures 2A & 2C show a phylogenetic tree representing the relationship among these cephalopods along with the percent of the genome occupied by each of four repeat element classes: DNA transposons, LINEs, LTRs, and SINEs. It is clear from this comparison that the lineage leading to octopuses underwent a dramatic increase in the number of SINE elements, with SINEs taking up approximately 4.64% of the genome sequence in *O. bimaculoides*. The SINE expansion in octopus genomes has been noted previously with sparser comparisons (Albertin et al., 2015), however when seen in a phylogenetic light (Fig. 2) this pattern is abundantly clear.

While a historical expansion of SINE elements is a striking feature of the octopus genome, we also found an abundance of LINE elements, which composed a higher proportion of the genome than SINE elements, at 12.73%. Two clades of LINE elements, R2/R4/NeSL and RTE/Bov-B, made up the bulk of this with 3.10% and 4.59% of the genomic sequence respectively (Fig. 2C).

Our chromosome-level assembly allowed us to explore if there is genomic heterogeneity in the accumulation of transposable elements (TEs). Visualization of the TE contributions to individual chromosomes (Figs. 2D, S3, S4, S5, S6) suggests that while most chromosomes have little variation in their proportions of TEs, the *O. bimaculoides* Z chromosome has an elevated density of LINE elements, harboring LINEs at approximately twice the density as the genome-wide background (Mann-Whitney test; *p*< 0.0001; Fig. 2D).

## LINE elements are a signature of coleoid sex chromosomes

We hypothesized that the abundance of LINE elements could be a clear signature of Z chromosomes in other cephalopod genome assemblies. Looking at the landscape of TEs in the high quality, chromosome-level assemblies of *O. sinensis, E. scolopes*, and *S. esculenta* yielded a striking pattern: a single large chromosome in each of those assemblies (chrZ in *S. esculenta*, formerly chr20 in *O. sinensis* and formerly chr43 in *E. scolopes*) contained a high density of LINE elements, perhaps suggesting the presence of an orthologous Z chromosome (Fig. 2D). As in *O. bimaculoides*, each of these chromosomes harbors a significantly greater density of LINEs in comparison to the rest of their genomes (Mann-Whitney tests; *p* < 0.0001, *p* = 0.00023 and *p* < 0.0001, for *O. sinensis, Euprymna scolopes*, and *Sepia esculenta* respectively; Table S5). As a result of these findings supporting our hypothesis, we renamed the putative Z chromosomes to Z in all figures for clarity. Upon inspection of the much more distant *Nautilus* genome, no chromosomes with elevated LINE element content could be identified. Thus evidence from LINE element enrichment suggests that this could be a unique feature of the coleoid Z chromosome that may have originated before the split of the squid and octopus lineages, but after divergence from *Nautilus*.

## Syntenic relationships of genes on the Z chromosome are conserved

We hypothesized that the chromosomes with elevated LINE densities in *O. sinensis, E. scolopes*, and *S. esculenta* might be orthologous to the *O. bimaculoides* Z chromosome. To test this hypothesis, we compared synteny among chromosomes using gene-level annotations as revealed using the GENESPACE and MCScanX software packages (Lovell et al., 2022; Wang et al., 2012, 2024), which use orthologous gene identification to define conserved synteny blocks. As seen in Figures 3A and 3B, gene-based synteny confirms our hypothesis – the *O. bimaculoides* Z chromosome has a large block of conserved synteny on the putative Z chromosomes in *O. sinensis, E. scolopes*, and *S. esculenta*. Dotplots showing finer resolution pairwise alignments are shown in Figure S8.

**Figure 3.**
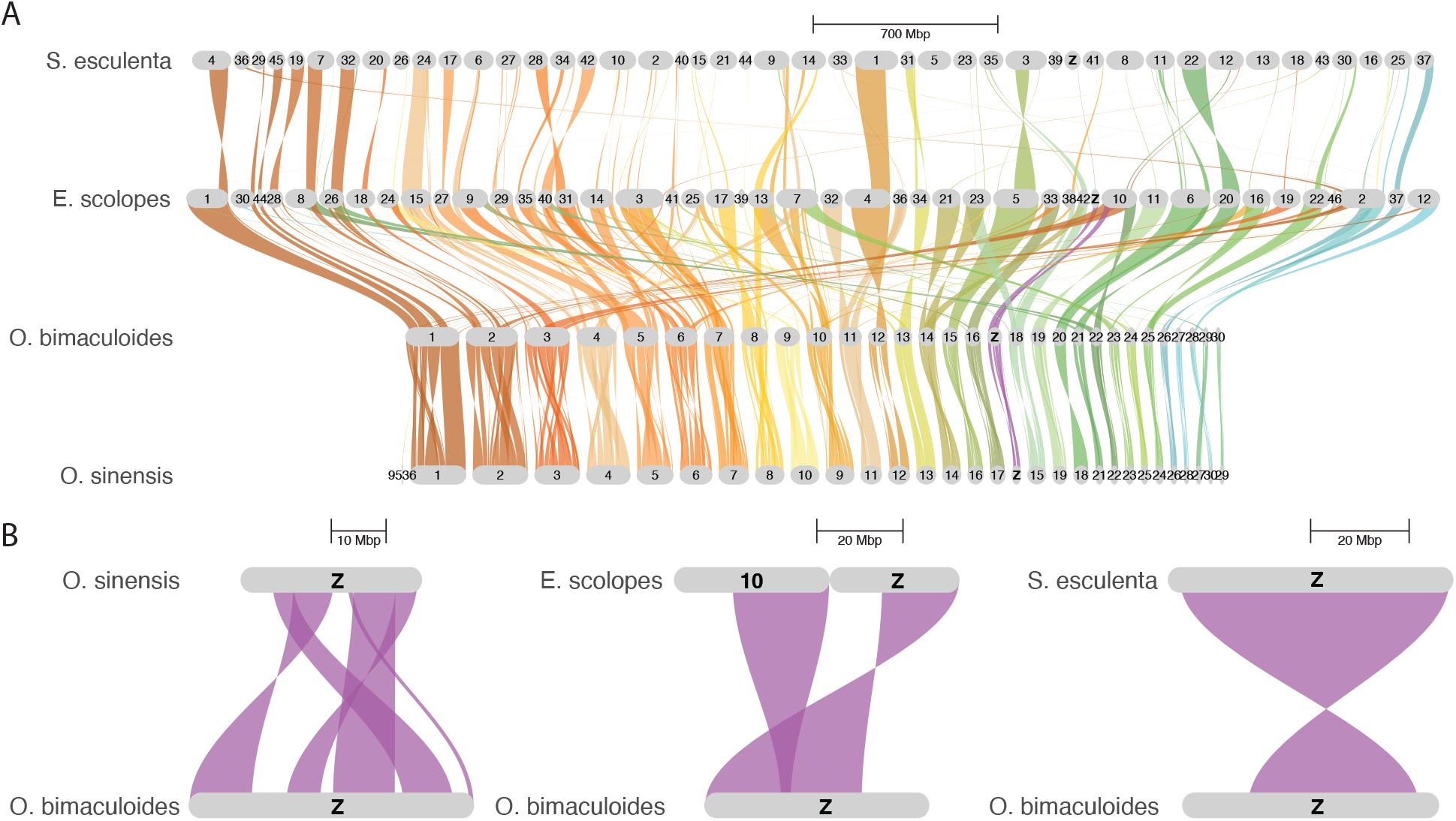
Syntenic relationships between four cephalopod species. **A**. Riparian synteny plot generated using orthogroups between the chromosomes of representative octopus, squid, and cuttlefish species. Ribbons connected to the *O*.*bimaculoides* Z chromosome are highlighted in purple. **B**. Paired riparian synteny plots between the Z chromosome of *O. bimaculoides* and putative Z chromosomes of three other species, *O. sinensis, E. scolopes*, and *S. esculenta*, reveal conservation of the Z chromosome.

Due to the huge evolutionary distances between *Nautilus pompilius* and the other cephalopod taxa studied here, coupled with the extensive genome expansion along the coleoid lineage, calling syntentic regions on the basis of colinear blocks of one-to-one orthologs across multiple taxa is challenging. Instead we used MCscanX, a method not constrained to singlecopy genes, to compare chromosome Z of *O. bimaculoides* to the *N. pompilius* genome in a more targeted fashion. This revealed 1,057 collinear genes (8.31% of all genes) contained on 135 alignment blocks between *O. bimaculoides* Z-linked proteins and *N. pompilius*. Syntenic blocks are found across almost all of the *N. pompilius* chromosomes, indicative of the extensive genome and gene duplication since coleoid cephlapods diverged from *N. pompilius* (Fig. S9). Of the 300 single copy, unique Z-linked genes in *O. bimaculoides*, more than half were contained on one of two chromosomes in *N. pompilius*: 87 were contained in collinear blocks on chromosome 7 and 77 were found on *N. pompilius* chromosome 4. We note that neither of these chromosomes is unusual for LINE density (see Fig. 2D).

## A single, ancient origin of the cephalopod Z chromosome

Our evidence, when taken together, demonstrates that the *O. bimaculoides* Z chromosome is a genomic outlier with clear homology at the gene level and with respect to repeat content to the putative Z chromosomes in *O. sinensis, E. scolopes*, and *S. esculenta*. While the divergence time between *O. bimaculoides* and *O. sinensis* is relatively modest, between 13 - 34 million years (Huang et al., 2022), divergence between Decapodiformes (squid and cuttlefish) and Octopodiformes (octopus) is much older, with an approximate divergence time between *O. bimaculoides* and *E. scolopes* of 285 - 340 million years (Huang et al., 2022). Thus, it was imperative to examine whether the orthologous chromosomes in other cephalopod species are hemizygous. Figure 4 presents sequencing coverage data from four additional species, including short read libraries that we generated from sexed *Octopus bimaculatus* individuals (see methods), the sister taxa to *O. bimaculoides*, a female cuttlefish *S. esculenta* recently released by the Darwin Tree of Life project (NCBI, GCA 964036315.1), Illumina short read libraries of two unsexed individuals of *N. pompilius* (Zhang et al., 2021; Huang et al., 2022) and a set of eight Illumina short read libraries derived from single, unsexed embryos of *E. scolopes* from Schmidbaur et al. (2022). These results again confirmed our hypothesis in each case. In species where we have sexed individuals (*O. bimaculatus* and *S. esculenta*), the putative Z chromosome appears at approximately half coverage in the female in comparison to representative autosomes (chromosomes 1 and 10 shown as autosomal comparisons in Figs. 4A, 4B; all chromosomes shown in S10, and S11). For *Euprymna scolopes* we did not have sexed individuals, however in the eight embryos coverage among chromosomes was nearly uniform across individuals except at the putative Z (chr43). The putative Z is an outlier in comparison to the other chromosomes with multiple distinct coverage classes among individuals, suggesting females are hemizygous as observed in *O. bimaculoides* (Figs. 4C; all chromosomes shown in S12, & S13). Finally, in comparing sequencing coverage among the two available unsexed *Nautilus* genomes, we again observed a single chromosome, chr4, at reduced coverage in one of the two libraries (Fig. 4D). Chr4 is one of two chromosomes in *Nautilus* that shows synteny to the *O. bimaculoides* Z as determined by our MCScanX analysis. Thus, while we were not able to detect a Z chromosome based on LINE element density in *Nautilus*, sequencing coverage among individuals and synteny strongly suggests that a sex chromosome is present and that it shows orthology to the Z in coleoid cephalopods.

**Figure 4.**
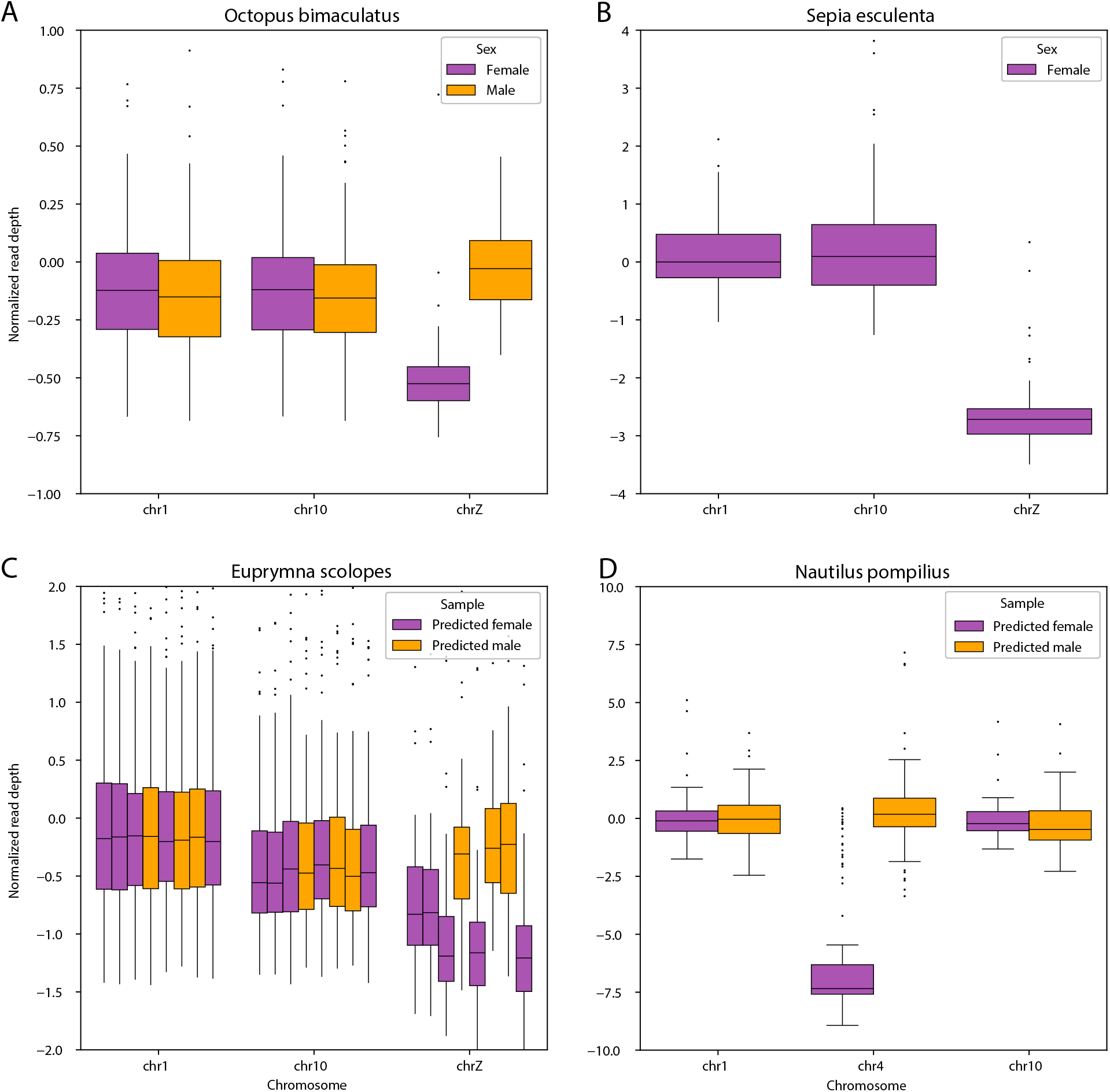
Normalized coverage of sequencing data mapped to representative chromosomes in *O. bimaculatus, S. esculenta, E. scolopes* and *N. pompillius*. **A**. For *O. bimaculatus*, we generated Illumina short reads from two male and two female individuals (See Methods for more details). **B**. Female *S. esculenta* short reads were generated by Darwin Tree of Life Project and downloaded from NCBI (GCA 964036315.1). **C**. Sequencing data for unsexed embryos of *E. scolopes* were generated by Schmidbaur et al. (2022). The putative Z chromosome is the only chromosome with variation in coverage across individuals (see Figs S12 & S13). Predicted female individuals (purple) have significantly lower coverage at the presumed Z chromosome compared to the average coverage of all other chromosomes (*p*< 0.0005 in all cases; Mann Whitney U tests) and predicted males (orange) do not (N.S.; Mann-Whitney U tests;). **D**. In *N. pompilius*, Illumina short reads from the predicted female individual were generated by Zhang et al. (2021) and Illumina short reads from the predicted male individual were generated by Huang et al. (2022).

Having established strong support for homology of the Z chromosome among cephalopods, we were lastly interested in examining what genes might be shared in orthologous regions of the Z chromosome among lineages. We first compared annotations from the shared syntenic chromosome Z blocks defined by GENESPACE across coleoid cephalopods, using annotation from *O. bimaculoides, O. sinensis*, and *E. scolopes*. This revealed 19 unique protein coding loci shared among these three taxa that are housed in this region of the genome. We did blastp homology searches of these genes to human proteins and found 16 of 19 had strong hits (Table S6). Using publicly available summaries from GeneCards (Stelzer et al., 2016) we report that all 16 show mRNA expression in human reproductive tissues, and 15 of 16 of these show protein expression in human reproductive tissues. A particularly leading hit is the protein obimac0008950, which shows strong homology to the human Sperm Associated Antigen 9 (SPAG9; e-val=0.0). We next asked if the 19 proteins we identified above also occur in the colinear blocks identified between the Z in *O. bimaculoides* and *N. pompilius* and found that 9/19 were present, and 6 of these 9 could be found in humans. All 6 show evidence of mRNA expression and 5 of these 6 show protein expression in human reproductive tissues. Taken together, these blast results let us speculate that the genes retained on the Z chromosome may represent an ancient set of proteins essential to animal reproduction and/or gametogenesis. Last, we searched for known sex determination genes from other mollusc species on the Octopus Z. Four are described: *Forkhead box protein L2* (*FOXL2*) in *M. yessoensis* and *C. farreri, Zinc finger protein 226-like* (*ZNK226l*) in *P. magellanicus*, and *Cytochrom P450 3A24-like* (*CYP3A24l*) in *A. japonicum* have been found to be female-biased (Han et al., 2022). *Doublesex and mab-3 related transcript factor 1-like* (*DMRT1L*) is mollusc-specific and male-biased in *M. edulis* (Evensen et al., 2022). We downloaded representative protein sequences of these genes from NCBI (*FOXL2* and *DMRT1L* from *A. irradians, ZNK226l* and *CYP3A24l* from *M. yessoensis*) and ran BlastP searches to a database composed of proteins we annotated on the *O. bimaculoides* Z chromosome. This revealed Z-linked hits for *FOXL2* (obimac 0009054.1, e-val=3.23e-33) and *ZNK226l* (obimac 0008819.2, e-val=2.66e-27), thus the *O. bimaculoides* Z harbors known mollusc sex determination genes.

Our results provide the first glimpse of sex determination in cephalopods, a phenomenon which until now has remained a mystery. The clear presence of a hemizygous Z chromosome in females suggests a ZZ/ZO sex determination system that had previously been missed in cytological comparisons. Since ovaries contain fewer cells in meiosis than testes, it is difficult to identify female heterogameity via karyotyping (Jonika et al., 2022). Therefore, it is standard to use testicular tissue (Bonnaud et al., 2004; Vitturi et al., 1982) or unsexed embryos (GAO and Natsukari, 1990) for karyotyping. Although we have not found evidence for a W chromosome in *O. bimaculoides* or any other cephalopod species, assembly of degenerate chromosomes is particularly challenging and so evidence for a W might be gathered in the future. Taken together, our evidence suggests that the cephalopod Z chromosome evolved once in the lineage leading to the common ancestor of all extant lineages. Recent molecular estimates of the time to the common ancestor of *Nautilus* and coleoids place it at ∼ 482 million years ago (range 398-520 Mya; Huang et al. (2022)) thus the Z chromosome would had to have originated at least this long ago. This is an astoundingly long time for a sex chromosome to be preserved (Charlesworth et al., 2005). A few other ancient sex chromosomes have been described previously from liverworts (∼ 430Mya; Iwasaki et al. (2021)) and mosses (∼ 300Mya; Carey et al. (2021)), and there is evidence that the insect X chromosome is quite ancient (∼ 450Mya; Toups and Vicoso (2023)), although it shows rapid evolution in some clades. Thus in context, the cephalopod Z chromosome is among the very oldest sex chromosomes yet described.

## Acknowledgments

We thank Caroline Albertin and Matthew Birk for providing samples, Mara Lawniczak for her input on the project and comments on the manuscript, and John Postlethwait, Peter Ralph, Melissa Toups, Graham Coop, Clara Rehmann, and members of the Kern Ralph colab for their input and comments on this manuscript. Sequencing and sample preparation was performed by the University of Oregon Genomics & Cell Characterization Core Facility. GCC was funded by NSF GRFP 1842486. ADK was funded in part by NIH awards R35148253 and R01HG010774. CMN was funded in part by NIH award R01NS118466. CMN and ACM were funded in part by a University of Oregon Renee James Seed Grant.

## Author Contributions

G.C.C. and A.D.K designed the study. J.O.S.C., D.M.P., and J.R.P. prepared samples for sequencing. G.C.C. assembled the genome, G.C.C and S.T. annotated the genome, S.T. conducted the repeat analysis and generated the multiple species alignment, G.C.C. and S.T.S. conducted the synteny analyses. A.D.K did homology searches. G.C.C, S.T., S.T.S, and A.D.K wrote the manuscript. A.C.M, C.M.N, and A.D.K. helped supervise the project. All authors read and approved the final manuscript.

## Data availability

The genome assembly and associated sequencing reads have been deposited to BioProject ID PRJNA1076263. Additional data have been deposited to a Zenodo repository at 10.5281/zenodo.13278524. Silhouettes of cephalopod taxa were obtained from PhyloPic (https://www.phylopic.org/).

