## Supplement for "Cephalopod Sex Determination and its Ancient Evolutionary Origin"

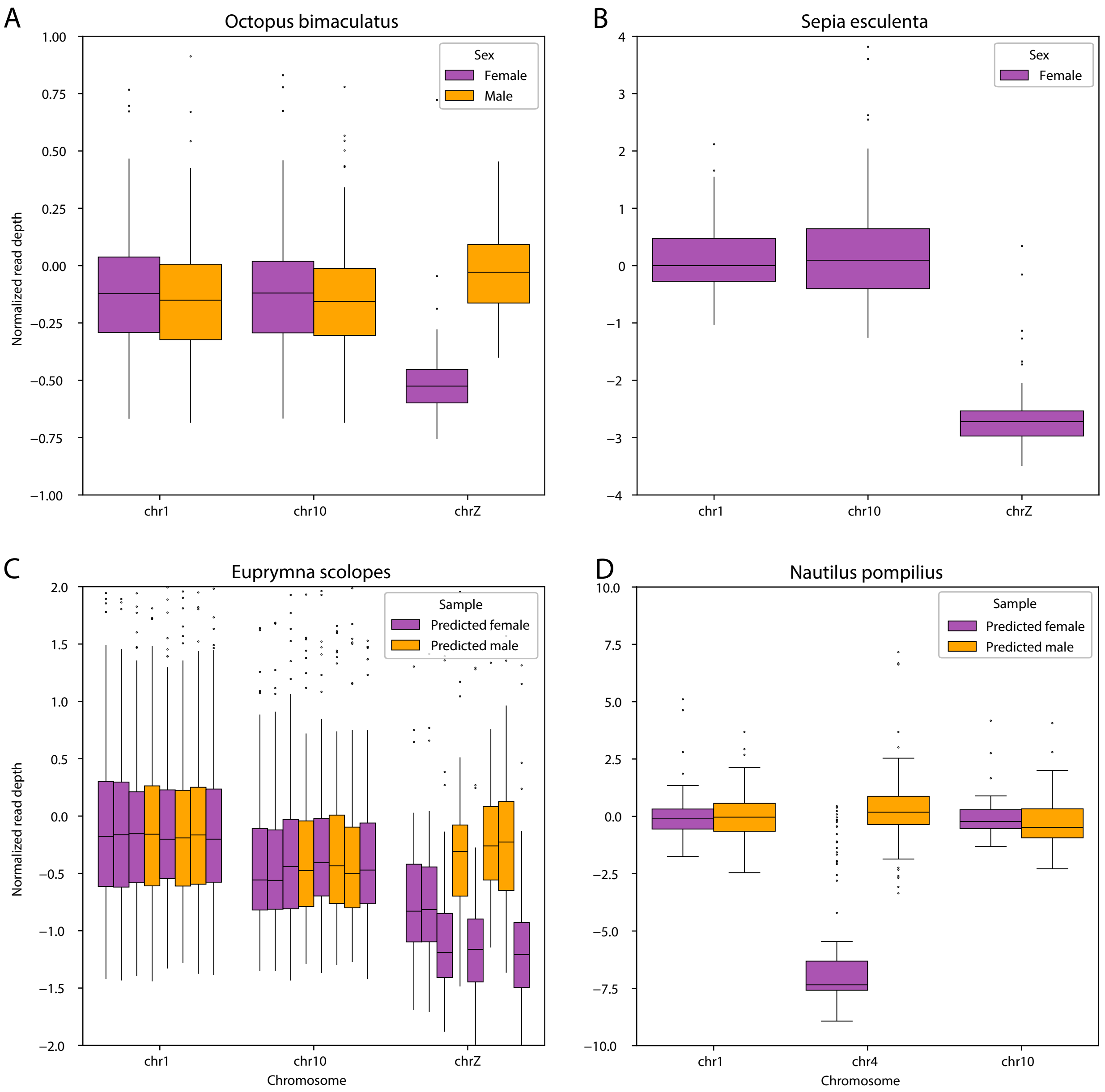


Figure 4: Normalized coverage of sequencing data mapped to representative chromosomes in *O. bimaculatus*, *S. esculenta*, *E. scolopes* and *N. pompillius*. **A.** For *O. bimaculatus*, we generated Illumina short reads from two male and two female individuals (See Methods for more details). **B.** Female *S. esculenta* short reads were generated by Darwin Tree of Life Project and downloaded from NCBI (GCA_964036315.1). **C.** Sequencing data for unsexed embryos of *E. scolopes* were generated by Schmidbaur et al. (2022). The putative Z chromosome is the only chromosome with variation in coverage across individuals (see Figs S12 & S13). Predicted female individuals (purple) have significantly lower coverage at the presumed Z chromosome compared to the average coverage of all other chromosomes ($p < 0.0005$ in all cases; Mann Whitney U tests) and predicted males (orange) do not (N.S.; Mann-Whitney U tests;). **D.** In *N. pompilius*, Illumina short reads from the predicted female individual were generated by Zhang et al. (2021) and Illumina short reads from the predicted male individual were generated by Huang et al. (2022).

### Supplemental Materials

#### Materials and Methods

#### Sample collection

A description of the genome sample collection and sequencing was published previously in Songco-Casey et al. (2022). Optic lobe tissue was collected from an adult female *O. bimaculoides* that was obtained from the southern coast of California. Optic lobe, central brain, retina, and arm tissue were dissected from related 6-week-old juvenile octopuses to be used for construction of an Iso-Seq library. Additional arm tissue was dissected to construct an Illumina library to be used in Hi-C sequencing. Four additional unrelated individuals were collected from the southern coast of California for sex chromosome coverage analysis. Optic lobe tissue was extracted from two individuals, gill tissue from one individual, and testes from one individual (Table S3).

#### Library construction and sequencing

All tissue samples were sent to the University of Oregon Genomics & Cell Characterization Core Facility (GC3F) for DNA extraction, sample preparation, and sequencing. High molecular weight genomic DNA was extracted with a Nanobind Tissue Big DNA kit (Circuloimics) from the adult optic lobe tissue. A SMRTbell Express Template Prep Kit 2.0 was used to construct a Pacific Bioscience standard HiFi library. Megaruptor 2 instrument was used to shear genomic DNA at 20kb target size and BluePippin size selection (Sage Science) omitted the smallest fragments (<10-14kb). Two HiFi genomic circular consensus sequencing (CCS) SMRTbell libraries were prepared as input for five HiFi SMRT cells. For Iso-Seq sequencing, a RNeasy Plus Mini Kit (QIAGEN) was used to conduct RNA extractions. GC3F prepared a multiplexed SMRTbell Iso-Seq library according to PacBio's protocol to be used as input for one SMRT cell. Single molecule sequencing of HiFi and Iso-Seq libraries was conducted with a Pacific Biosciences Sequel II system (Table S3). For scaffolding, we constructed one Illumina library with a Phase Hi-C kit to produce two NovaSeq 6000 sequencing Hi-C runs (Table S2). For the male vs. female coverage analysis, genomic DNA from each individual was barcoded with GC3F's TruSeq-style adaptors. The barcoded DNA was pooled and sequenced on two Illumina NovaSeq 6000 S4 lanes.

Post sequencing, the genomic HiFi reads were processed with SMRT Link to generate 5.8 million HiFi reads. The IsoSeq subreads were processed with the IsoSeq3 v.3.0.8 pipeline. Briefly, CCS generated HiFi reads. Primers were removed and barcodes were identified with lima. Reads were further refined by trimming poly(A) tails and removing concatemers. Finally, reads were clustered and collapsed into unique isoforms with parameters –min-aln-coverage 0.75 –min-aln-identity 0.75.

#### Genome size estimation and assembly

The *O. bimaculoides* genomic HiFi reads were used to estimate genome size. We generated a k-mer frequency distribution with JellyFish v.2.3.0 (Marçais and Kingsford, 2011), which was input into GenomeScope v.2.0 (Ranallo-Benavidez et al., 2020) to generate genome size estimation plots. Estimation with 21-mers estimated the genome to be 2.0Gb with 0.838% heterozygosity. Estimation with 31-mers estimated the genome to be 2.03Gb with 0.653% heterozygosity (Fig. S14).

Initial genome assembly was conducted using HiFiasm v.0.15.5-r352 (Cheng et al., 2021) with HiFi reads and standard parameters. Hifiasm did not sufficiently remove duplicates, so purge_dups v.1.2.5 (Guan et al., 2020) was used to purge haplotigs and overlaps. We placed the resulting scaffolds into a chromosome-level assembly using the data generated from the Hi-C library. The SLURM-compatible scripts of Juicer v.1.6 (Durand et al., 2016) were used to identify Hi-C contact points on the genome. 3D de novo assembly (3D-DNA) pipeline v.190716 (Dudchenko et al., 2017) was used to correct genome mis-assemblies, anchor order of chromosomes, and orient genomic fragments.

### Genome annotation

Transposable elements were identified using Repeat Modeler (Flynn et al., 2020) and were annotated with Repeat Masker v.4.1.5 (Chen, 2004). Protein-coding genes were annotated using both existing RNA-seq data and our newly generated Iso-seq data. Using existing RNA-seq databases, we ran Braker v.2.1.6 (Bruna et al., 2021) to annotate. Separately, we generated annotations with the Iso-seq reads using SQANTI3 v.5.1 (Tardaguila et al., 2018). The full-length isoforms generated by IsoSeq3 were filtered with SQANTI3 filter using default parameters. After generating the RNA-seq and Iso-seq based annotation files, they were merged using TAMA merge with parameters -e longest_ends -d merge_dup. Genome annotation statistics that were calculated with AGAT's agat_sq_stat_basic.pl script (Dainat, 2024) are presented in Table S2.

### Orthology inference

OrthoFinder v.2.5.4 was used with default parameters to cluster the protein sequences of *O. bimaculoides* into ortholog groups using sequences from seven additional cephalopod species including *Hapalochlaena maculosa*, *Octopus minor*, *Octopus sinensis*, *Architeuthis dux*, *Euprymna scolopes*, *Sepia pharaonis*, and *Nautilus pompilius*. The rooted species tree was used as input for the multiple species genome alignment.

### Multiple species alignment

We constructed whole genome alignments of eight cephalopod species with available reference genomes (Table S5) using progressive cactus (v2.3.0 via Docker container) (Armstrong et al., 2020). We used the whole genome alignments and phastCons v.1.5 (Hubisz et al., 2011) to identify highly conserved elements on each scaffold. This alignment, as well as our new reference genome and annotation are available as a UCSC genome assembly hub from http://poppy.uoregon.edu/~ssmall/ucsc_genome_hubs/v2.1.0/Octo.v2.1.0/hub.txt

#### Coverage calculations

To estimate read coverage of each species, we aligned short reads with bwa-mem2 (Vasimuddin et al., 2019) and long reads with minimap2 (Li, 2018) to the respective chromosome-scale genomes. Sequencing depth was estimated using mosdepth (Pedersen and Quinlan, 2018) in windows of 500,000bp. Plotted coverages of all species were normalized by the median coverage of the first chromosome.

#### Annotation of *Sepia esculenta*

At the time of these analyses, a genome annotation of *Sepia esculenta* was not yet publicly available. In order to include *Sepia esculenta* into the synteny analysis we lifted over the genome annotation from *Sepia pharaonis* (Song et al., 2021) onto *Sepia esculenta* using the software package Liftoff with default settings (Shumate and Salzberg, 2021).

#### Chromosome renaming

To simplify our analyses and plots we renamed all full-length chromosomes in all of the genomes that we included in this manuscript. We renamed chromosomes from longest to shortest with the naming scheme chr1, chr2, chr3, etc. with the exception of putative sex chromosomes which were renamed as chrZ.

#### Assessing sequence divergence among scaffolds

We converted the hierarchical alignment (hal) generated by progressive cactus to maf with hal2maf. We filtered the full, repeat-masked alignment into 1Mb windows along the genome using the phast utility msa_view, where windows were made with bedtools makewindows v2.31.0 (Quinlan and Hall, 2010). We further filtered alignments with multiple sequences per species using maftools mafDuplicateFilter v.0.1 (Mayakonda et al., 2018). We used phyloFit to construct phylogenies on each resulting 1Mb alignment region, only using alignments that had less than 40% of positions with gaps using the GTR mutation model. We calculated divergence as the cophenetic distance between *O. sinensis* and *O. bimaculoides* using the cophenetic.phylo function from the R package ape v.5.7-1 (Paradis and Schliep, 2019). The ratio of male to female mutation rates, $\alpha$, was estimated according to (Miyata et al., 1987). Let $R$ represent the ratio of Z-linked divergence to autosomal divergence. Then $\alpha = \frac{3R-2}{4-3R}$. $\alpha$ was estimated separately for mean dS at the invidual gene level as well as for mean divergence in 1Mb genomic windows. Confidence intervals on $\alpha$ were computed using bootstrap resampling of the observed data.

#### Synteny analyses

Synteny analyses between all chromosomes of *O. bimaculoides*, *O. sinensis*, *E. scolopes* and *S. esculenta* were conducted using the R package GENESPACE v.1.2.3 (Lovell et al., 2022). *O. bimaculoides* was used as the reference species for all synteny maps. In all riparian plots and dotplots, chromosomes are scaled by physical position (useOrder = FALSE parameter in GENESPACE).

To identify the macrosynteny relationships between *N pompilius* and other cephalopod species, we performed a macrosynteny analysis using MCScanX (Wang et al., 2012). Input files were prepared following protocols described in (Wang et al., 2024) and using protein sequences assigned to the Z chromosome in *O. bimaculoides* and 15,086 *N pompilius* proteins assigned to chromosomes (Huang et al., 2022). MCScanX was then run with default parameters and results visualized using scripts included with MCScanX program (www.github.com/wyp1125/MCScanX).

### dN and dS estimations

To estimate dN and dS, we used protein alignments of *O. bimaculoides* and *O. sinensis.* Proteins of 8,245 single copy orthologs previously identified using OrthoFinder v2.5.4 (Emms and Kelly, 2019), were re-aligned using MACSE v2.07 (Ranwez et al., 2021), with -prog alignSequences and default options. Z and autosome alignments were separated based on previous synteny analyses. In *O. sinensis*, chr20 (original name NC043016) was found to be orthologous to *O. bimaculoides* chrZ. Therefore, only proteins that were single-copy orthologs between the *O. bimaculoides* chrZ and the inferred *O. sinensis* chrZ were considered for the sex chromosome analysis. All other proteins were classified as autosomal. The module cal_dn_ds from BioPython (Cock et al., 2009) was used to estimate the non-synonymous and synonymous substitution in the alignment with options 'method=NG86'.

### GO term enrichment analysis

We conducted a GO term enrichment analysis of *O. bimaculoides* Z genes with the entire genome as the background with DAVID (Huang et al. (2009)). No functional clusters were found to be significantly enriched.

### Accession numbers for data used in this study

Accession numbers, sources, and citations for all data used and generated in this study are given in Table S7.

621 # Supplemental Figures

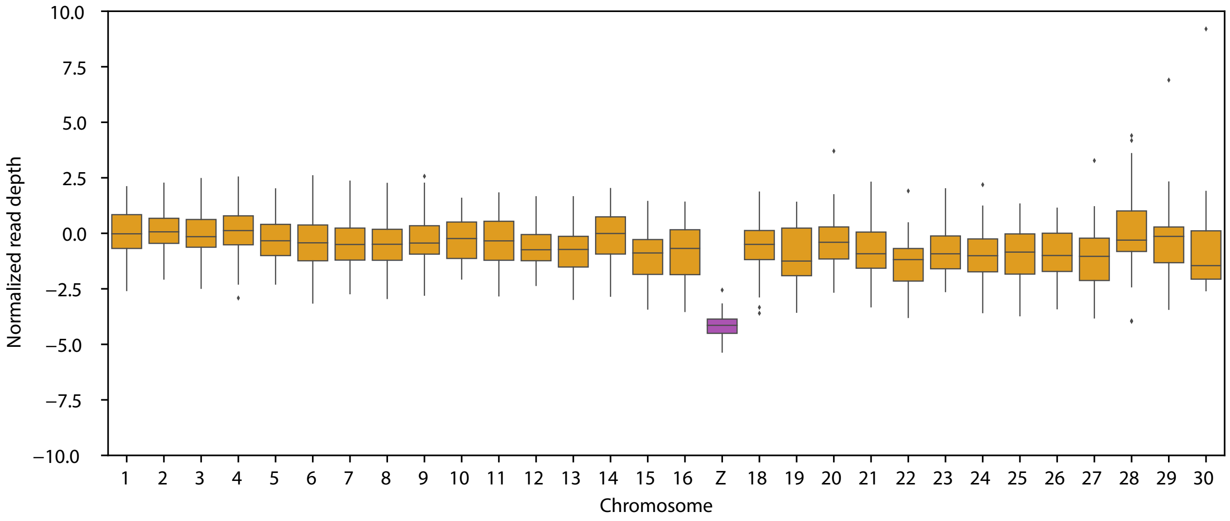


Figure S1: Normalized read depth of PacBio HiFi sequencing data across chromosomes from a female *O. bimaculoides.*

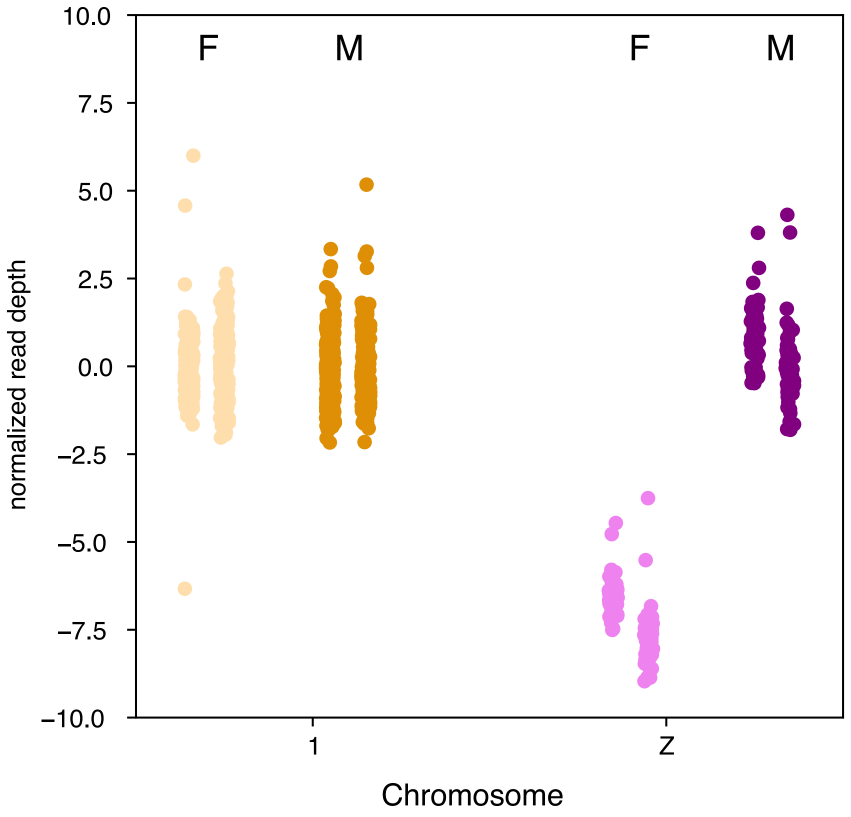


Figure S2: Normalized coverage of known sex individuals of *O. bimaculoides*. Includes two male (darker shades; labelled 'M') and two female (lighter shades; labelled 'F') with short-read sequencing. *O. bimaculoides* individuals at chromosomes 1 and Z are colored to reflect designation in main text. Both female individuals show hemizygous coverage at chromosome Z, whereas both male individuals show the same dosage at chromosome Z as at chromosome 1. See Table S3 for sequencing library information.

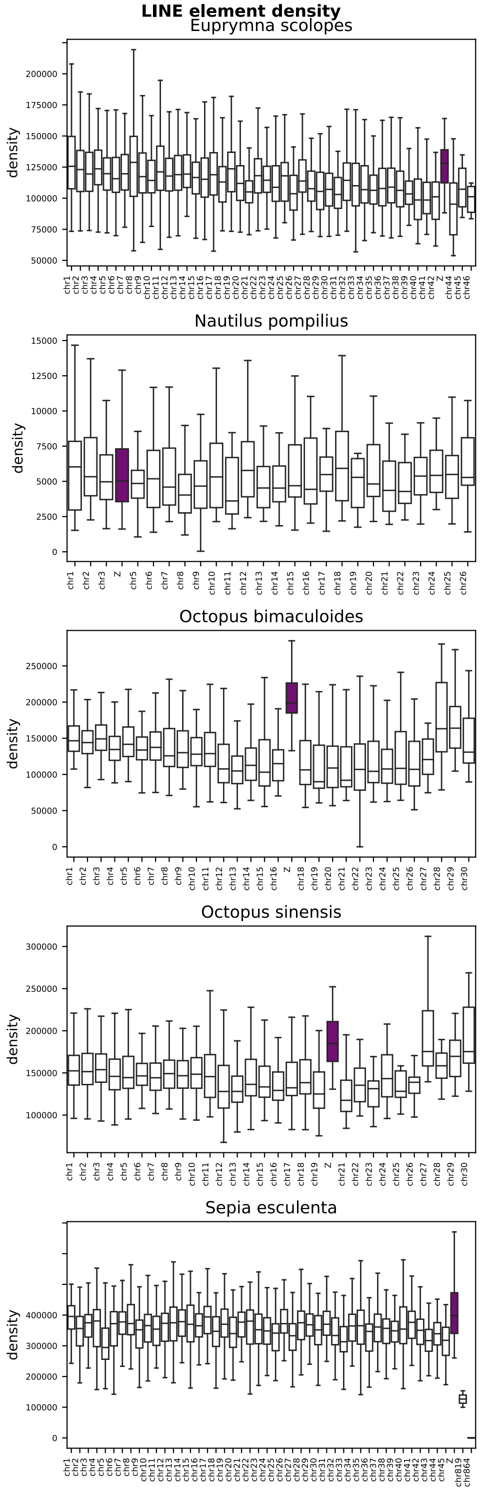


Figure S3: Density of LINE elements content on chromosomes arms across cephalopods. Data as in Figure 2 but with inclusion of *Nautilus*.

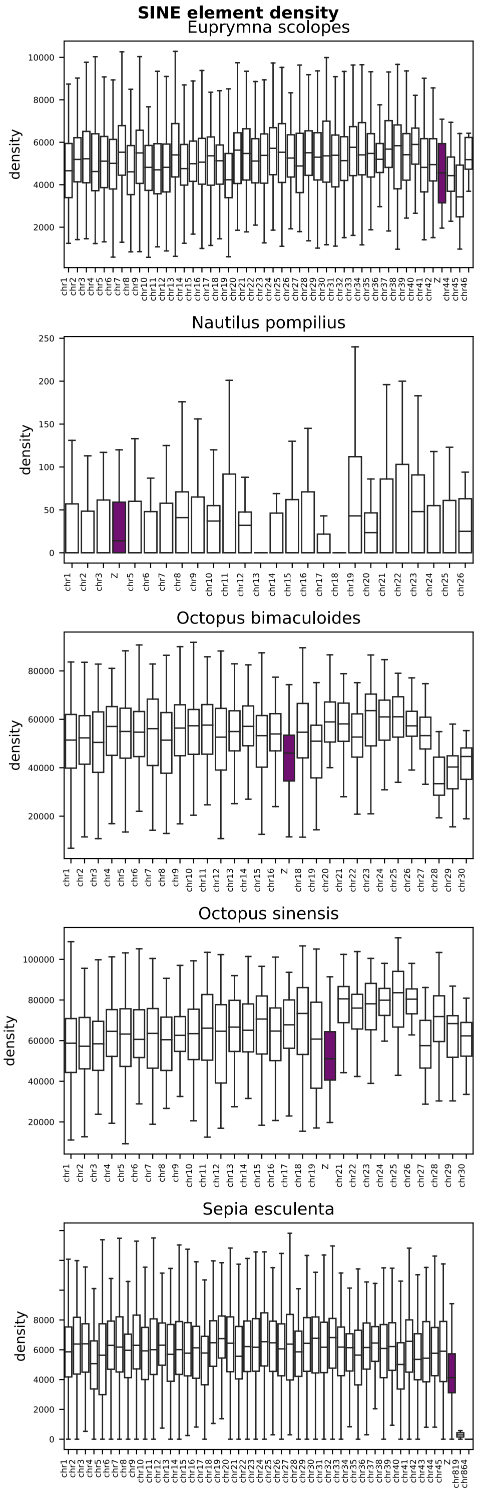


Figure S4: Density of SINE elements content on chromosomes arms across cephalopods.

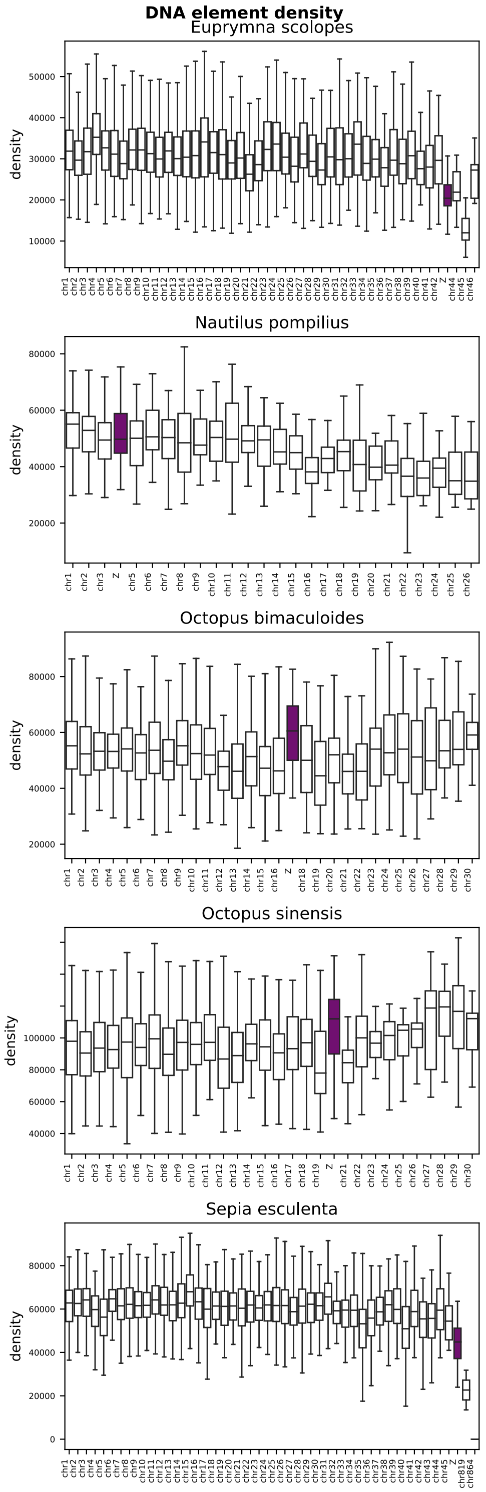


Figure S5: Density of DNA elements content on chromosomes arms across cephalopods.

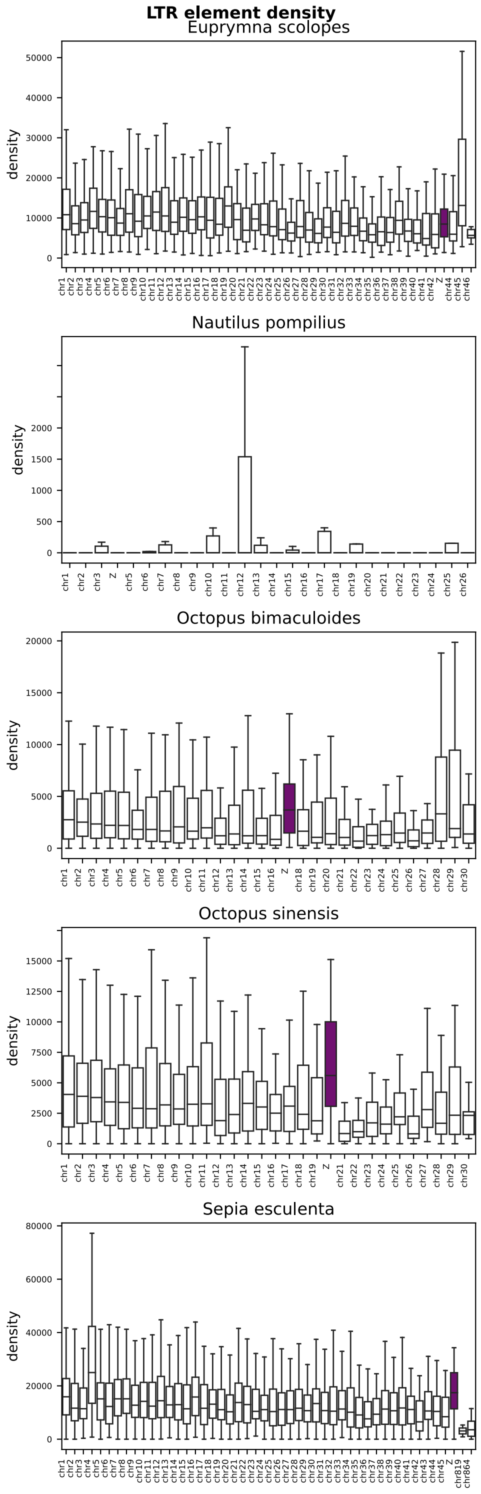


Figure S6: Density of LTR elements content on chromosomes arms across Cephalopods.

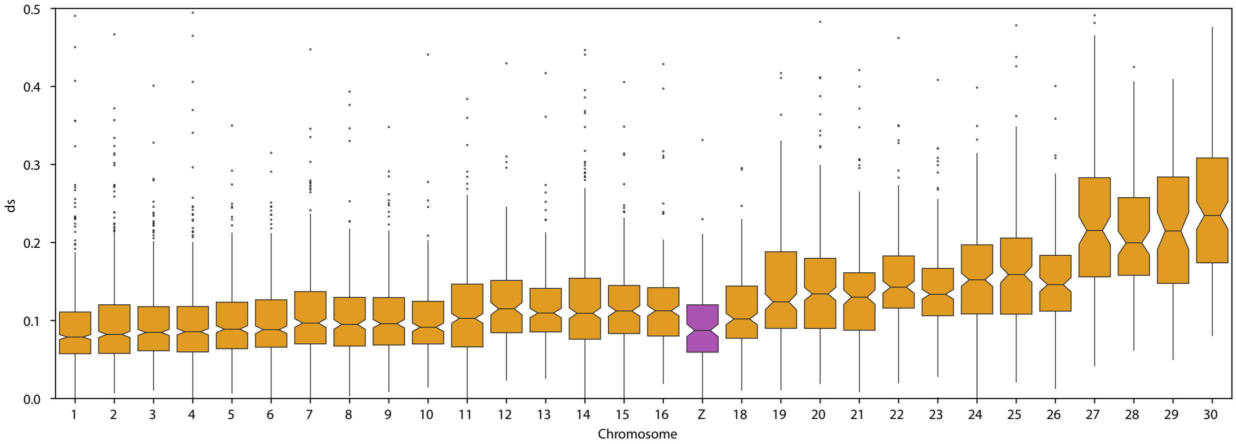


Figure S7: Synonymous substitution rate (dS) between *O. bimaculoides* and *O. sinensis* calculated for each gene per chromosome arm. The Z chromosome is highlighted in purple whereas autosomes are colored orange.

| | **Albertin et al. 2015** | **Albertin et al. 2022** | **This Assembly** |
|---|---|---|---|
| **Size** | 2.34 Gb | 2.37 Gb | 2.34 Gb |
| **Number of scaffolds** | 151,674 | 145,326 | 583 |
| **Average contig/scaffold length** | $15,415$ bp | $6,399$ bp | 4.02 Mb |
| **Longest scaffold** | 4.06 Mb | 199.87 Mb | 202.95 Mb |
| **Contig N50** | $5,532$ bp ($n = 97,860$) | $5,420$ bp ($n = 100,821$) | 0.86 Mb ($n = 775$) |
| **Scaffold N50** | 0.475 Mb ($n = 1333$) | 96.88 Mb ($n = 9$) | 101.05 Mb ($n = 8$) |
| **Percent gaps** | 15.134% | 15.016% | 0.118% |
| **GC content** | 36.04% | 36.03% | 35.79% |
| **Coverage** | $\sim 60x$ | $\sim 60x$ | $34x$ |
| **BUSCO (eukaryote)** | C:91.8% [S:90.6%,D:1.2%], F:6.7%, M:1.5%, $n$ : 255 | C:92.6% [S:91.4%,D:1.2%], F:6.3%, M:1.1%, $n$ : 255 | C:92.2% [S:91.0%,D:1.2%], F:6.7%, M:1.1%, $n$ : 255 |
| **BUSCO (metazoa)** | C:94.1% [S:93.8%,D:0.3%], F:3.6%, M:2.3%, $n$ : 954 | C:94.5% [S:94.1%,D:0.4%], F:3.2%, M:2.3%, $n$ : 954 | C:93.9% [S:93.2%,D:0.7%], F:2.6%, M:3.5%, $n$ : 954 |

Table S1: Comparison of Genome Assembly Statistics. Comparison of the initial genome assembly Albertin et al. (2015), a subsequent reassembly of those same sequencing data Albertin et al. (2022b) and the present assembly we introduce here.

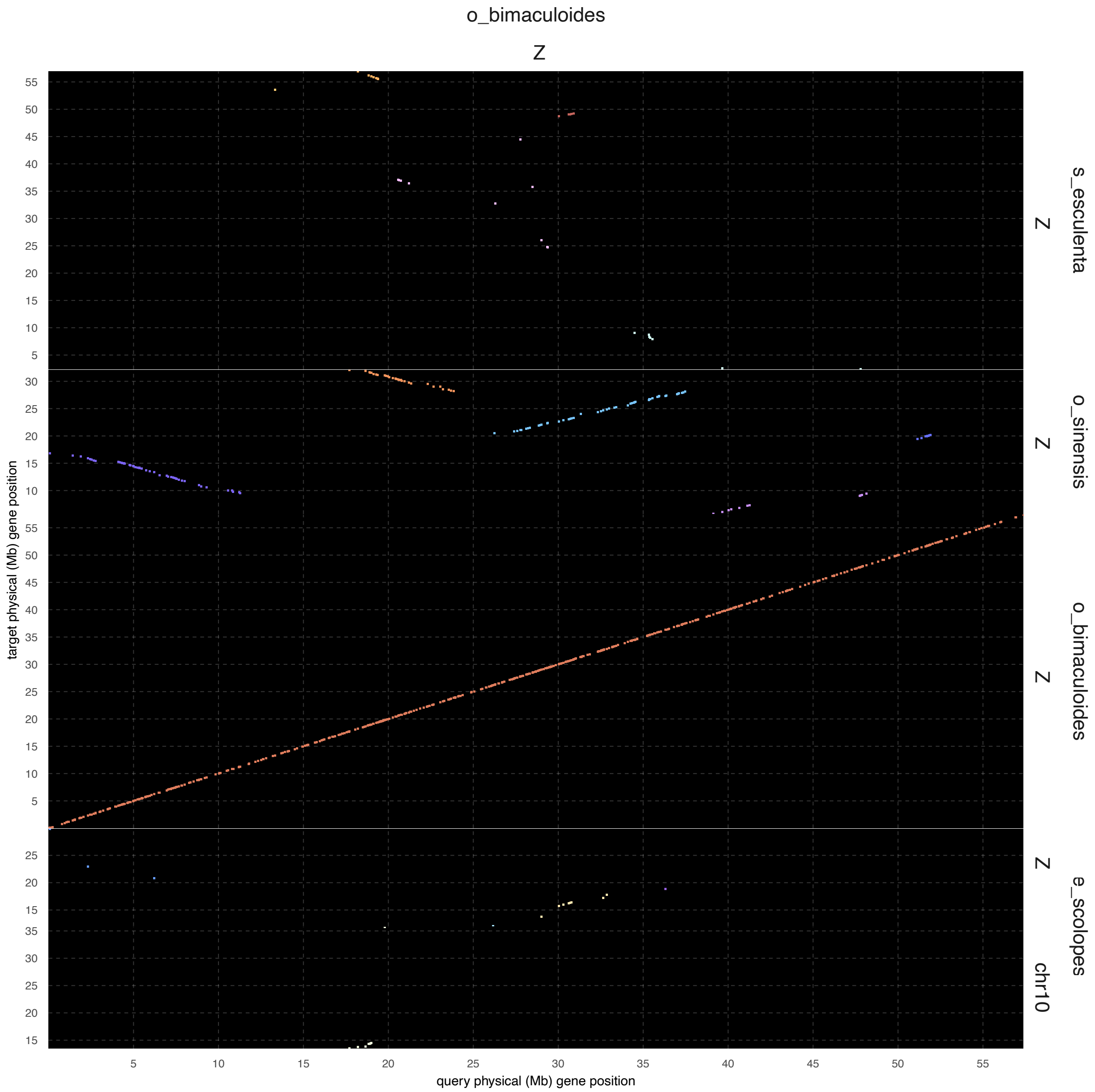


Figure S8: Dotplot of syntenic hits between the Z chromosomes of *O. bimaculoides*, *O. sinensis*, *E. scolopes*, and *S. esculenta*. The dotplot was generated with GENESPACE v.1.2.3 (Lovell et al. (2022)) and uses *O. bimaculoides* as the reference species.

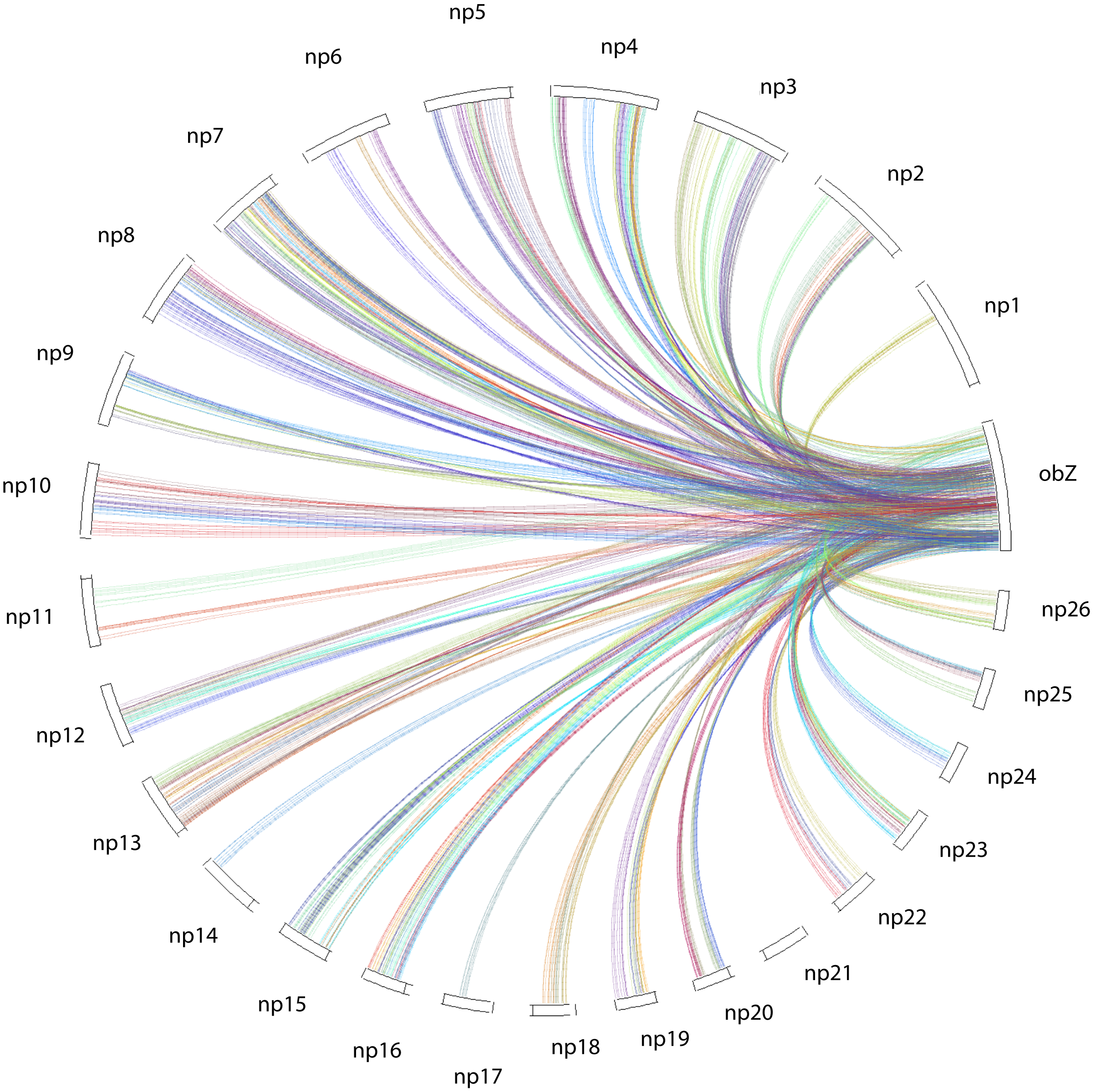


Figure S9: Syntenic blocks between the genes of *O. bimaculoides* Z chromosome and all genes of *N. pompilius*.

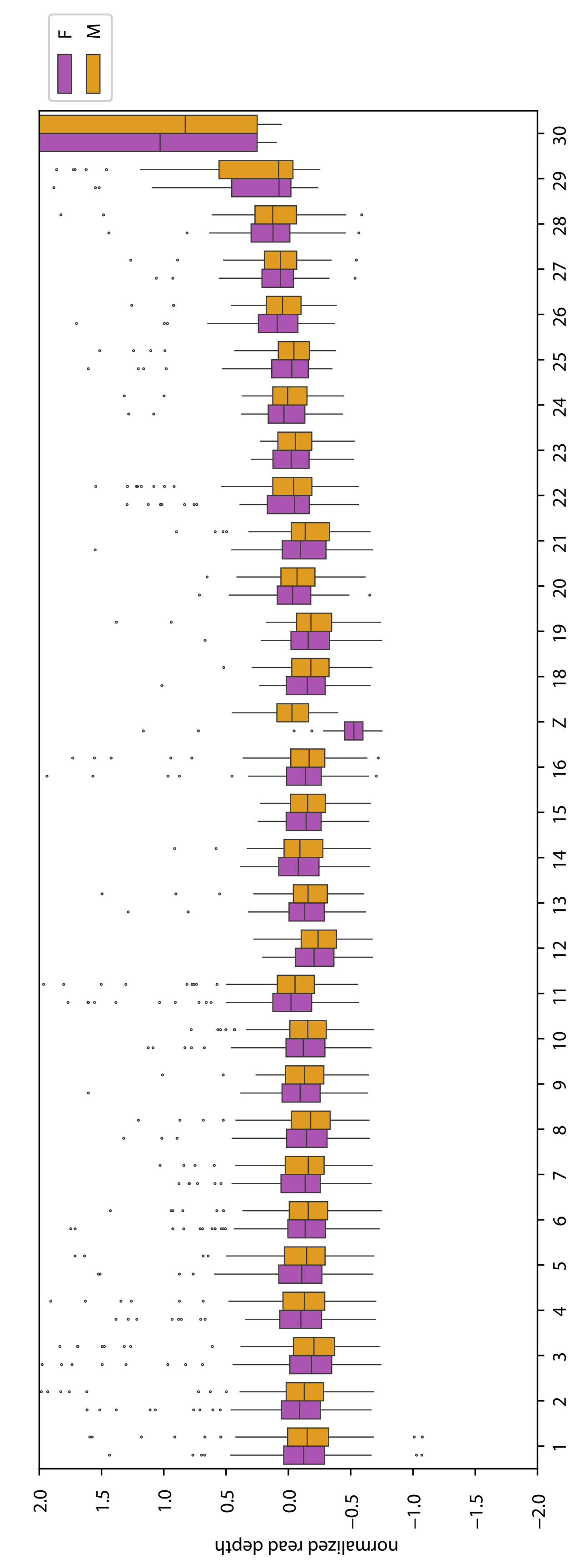


Figure S10: Normalized coverage of *O. bimaculatus* male and female short read libraries mapped to our *O. bimaculoides* genome assembly. Females have lower coverage than males in the Z chromosome.

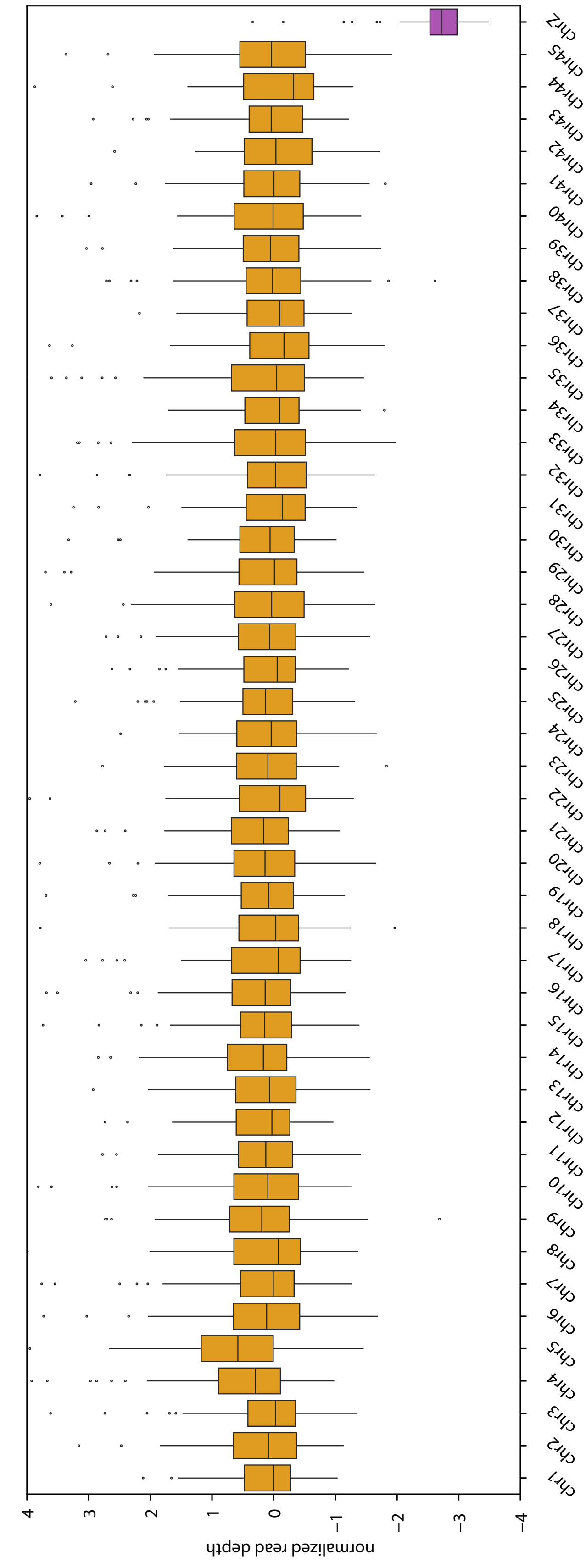


Figure S11: Normalized coverage of *S. esculenta* female short read library mapped to the *S. esculenta* genome assembly. Data and assembly was generated by the Darwin Tree of Life Project.

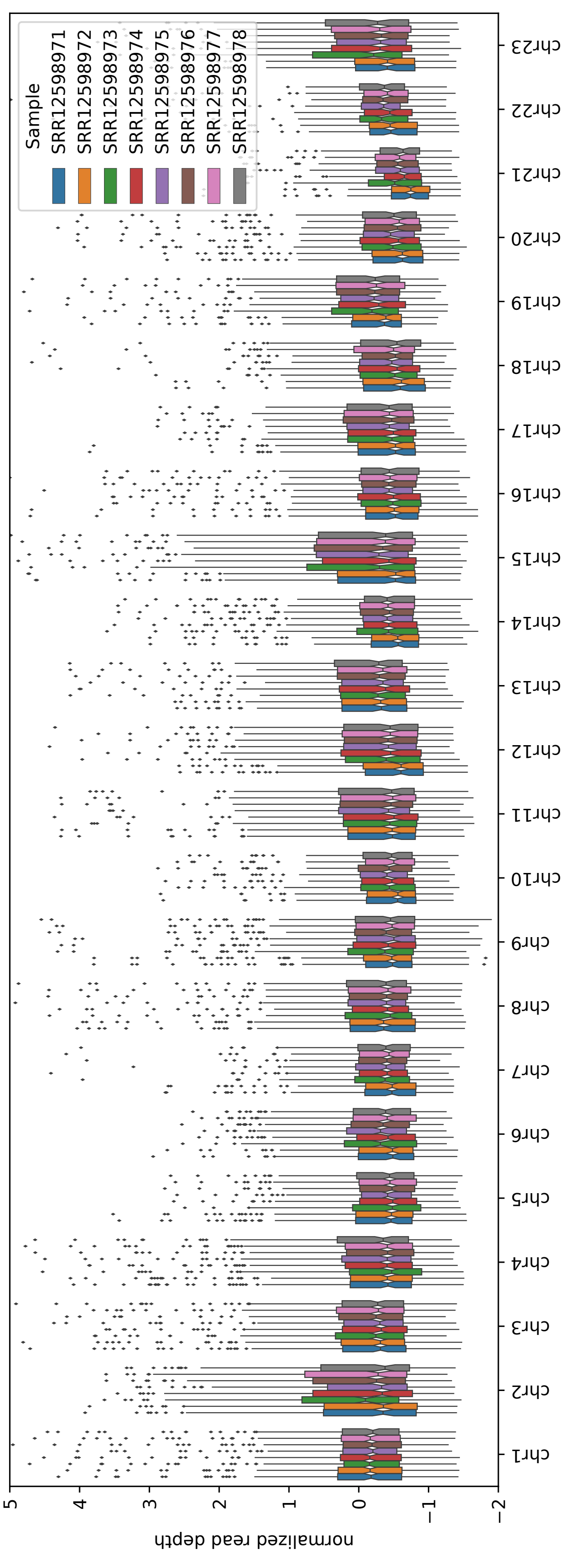


Figure S12: Normalized coverage of unsexed embryo short read libraries mapped to chromosomes 1 through 23 in *E. scolopes*. Chromosome 43 is the only chromosome with distinct variation in coverage. SRA identifiers for each library are shown in the legend. Data was generated by Schmidbaur et al. (2022).

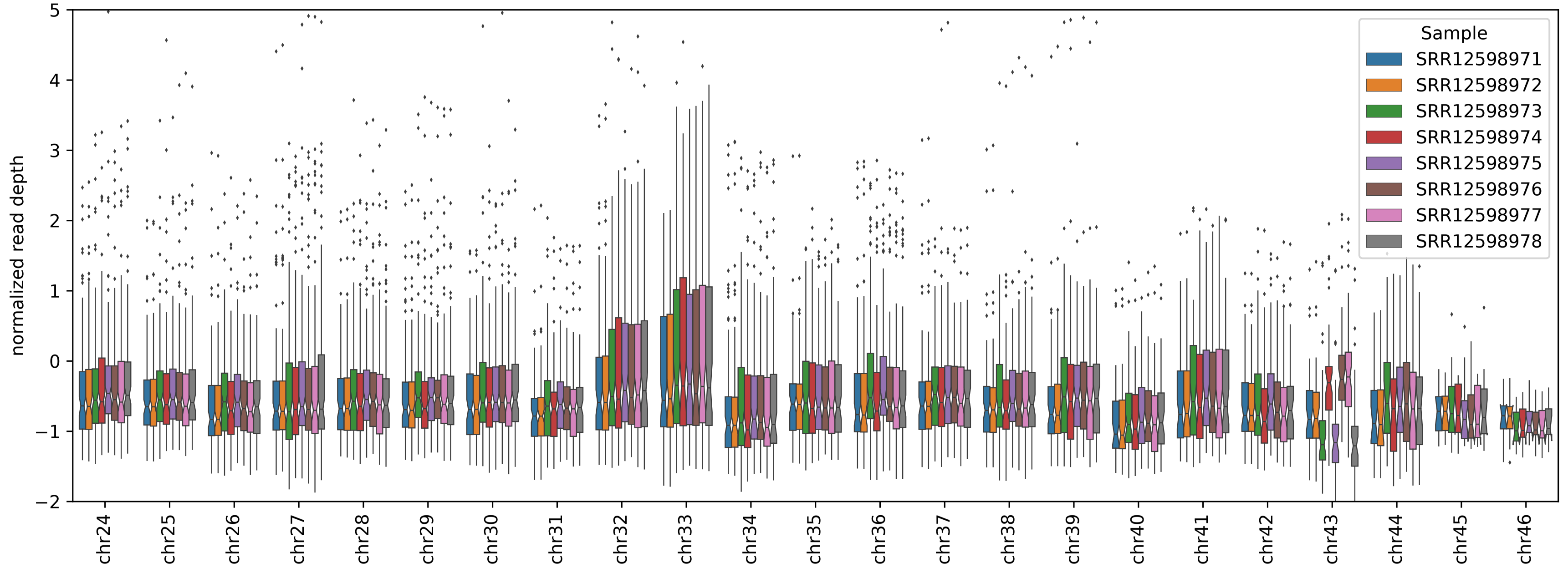


Figure S13: Normalized coverage of unsexed embryo short read libraries mapped to chromosomes 24 through 46 in *E. scolopes*. Chromosome 43 is the only chromosome with distinct variation in coverage. SRA identifiers for each library are shown in the legend. Data was generated by Schmidbaur et al. (2022).

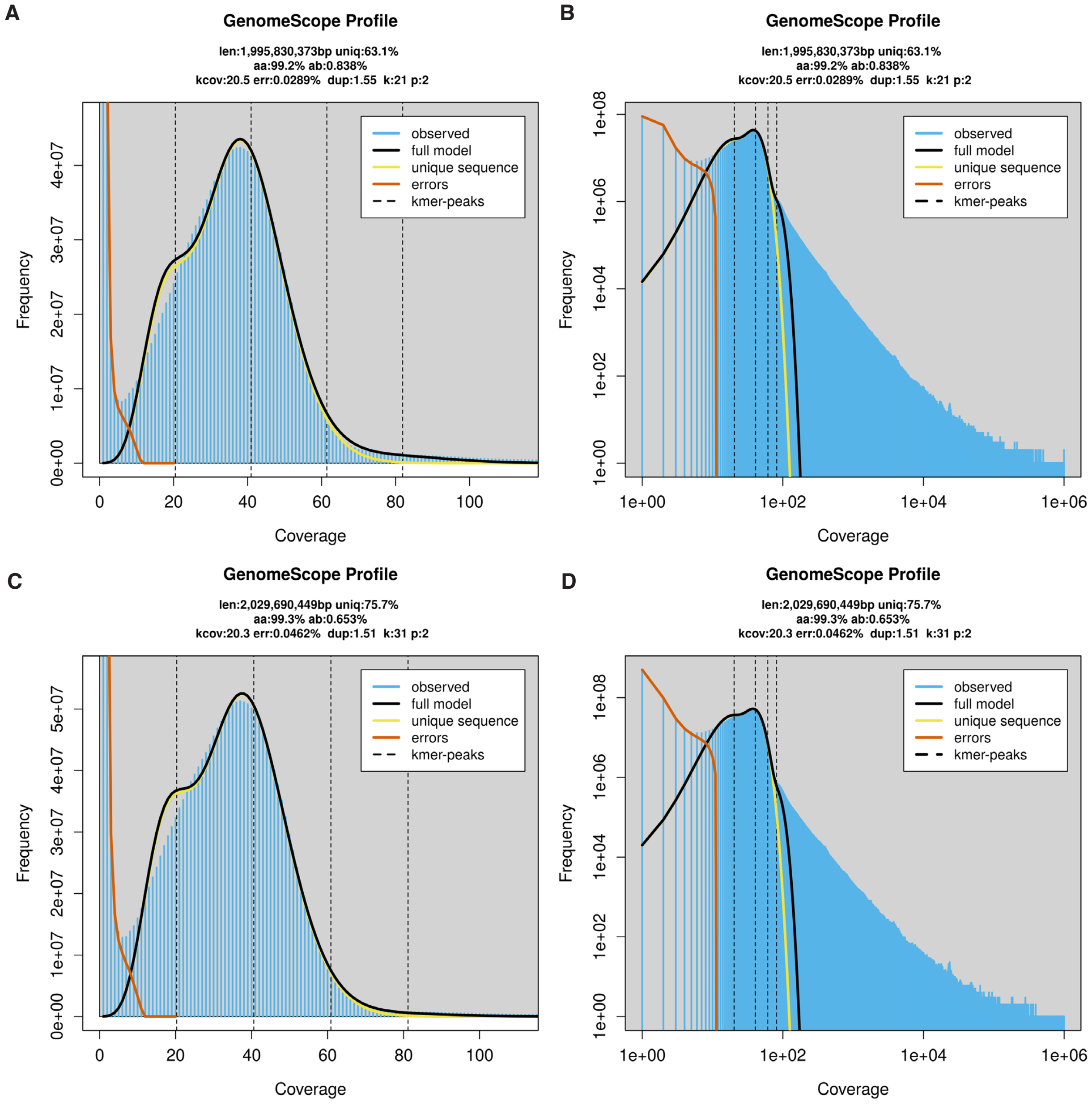


Figure S14: K-mer based genome size prediction of O. bimaculoides. **A** and **B** are 21-mer. **C** and **D** are 31-mer.

| | |
|---|---|
| **Number of genes** | 30655 |
| **Number of transcripts** | 61652 |
| **Number of cds** | 61639 |
| **% of genome is cds** | 1.3% |
| **Number of exons** | 504491 |
| **Mean transcripts per gene** | 2.0 |
| **Mean exons per transcript** | 8.2 |
| **Total gene length (bp)** | 758146852 |
| **Total transcript length (bp)** | 2799696948 |
| **Total cds length (bp)** | 77744706 |
| **Mean gene length (bp)** | 24731 |
| **Mean transcript length (bp)** | 45411 |
| **Mean cds length (bp)** | 1261 |
| **Mean exon length (bp)** | 279 |
| **Mean intron in cds length (bp)** | 5154 |
| **Mean intron in exon length (bp)** | 6004 |

Table S2: *O. bimaculoides* genome annotation statistics.

| Individual | Sex | Tissue | Data type | Total bases | Read Number | Depth | Notes |
|---|---|---|---|---|---|---|---|
| **Ob1** | F | optic lobe | HiFi, PacBio Sequel II 8M | 133.4Gb, post-CCS | 5.9M | 34x | five SMRT cells, two libraries |
| **Ob2** | hatchling | optic lobe, central brain, arm crown, retina | IsoSeq, PacBio Sequel II 8M | 10.4Gb | 1.6M (64,190 post IsoSeq3) | 0.04x | one SMRT cell |
| **Ob3** | hatchling | arm crown | Hi-C, Illumina NovaSeq 6000 | 555.8Gb | 3.7B | – | one flow cell, two runs |
| **Obmac_F_2** | F | optic lobe | Illumina NovaSeq 6000 | 90.3Gb | 1.24B | 71x | – |
| **Obmac_M_3** | M | optic lobe | Illumina NovaSeq 6000 | 76.2Gb | 1.05B | 60x | – |
| **Obmac_F_10** | F | gill | Illumina NovaSeq 6000 | 80.7Gb | 1.13B | 64x | – |
| **Obmac_M_14** | M | testes | Illumina NovaSeq 6000 | 66.0Gb | 924M | 52x | – |

Table S3: Stats of raw sequencing data collected from *O. bimaculoides* samples.

| **Individual** | **Sex** | **Tissue** | **Data type** | **Total bases** | **Number of reads** | **Depth** |
|---|---|---|---|---|---|---|
| **Oblat_F_2** | F | optic lobe | Illumina NovaSeq 6000 | 94.0Gb | 669M | 35x |
| **Oblat_M_3** | M | optic lobe | Illumina NovaSeq 6000 | 92.6Gb | 664M | 34x |
| **Oblat_F_10** | F | optic lobe | Illumina NovaSeq 6000 | 111.4Gb | 764M | 42x |
| **Oblat_M_4** | M | optic lobe | Illumina NovaSeq 6000 | 111.6Gb | 779M | 42x |

Table S4: Stats of raw sequencing data collected from *O. bimaculatus* samples.

| Species | Repeat type | p-value |
|---|---|---|
| *Sepia esculenta* | DNA | **4.06E-20** |
| *Euprymna scolopes* | DNA | **2.91E-17** |
| *Octopus bimaculoides* | DNA | **1.34E-04** |
| *Octopus sinensis* | DNA | 3.04E-03 |
| *Octopus bimaculoides* | LINE | **9.90E-25** |
| *Octopus sinensis* | LINE | **5.71E-09** |
| *Sepia esculenta* | LINE | **6.21E-05** |
| *Euprymna scolopes* | LINE | **2.34E-04** |
| *Octopus bimaculoides* | LTR | **1.23E-04** |
| *Octopus sinensis* | LTR | 1.06E-03 |
| *Sepia esculenta* | LTR | **3.64E-03** |
| *Euprymna scolopes* | LTR | 3.05E-01 |
| *Octopus bimaculoides* | SINE | **3.66E-06** |
| *Sepia esculenta* | SINE | **7.28E-06** |
| *Octopus sinensis* | SINE | 1.45E-03 |
| *Euprymna scolopes* | SINE | 1.80E-02 |

Table S5: Mann–Whitney U tests by repeat element type, testing for a significant difference between the sex chromosome element density against all autosomes. Bold indicates significance after Bonferroni correction.

| octo_gene_id | human_gene_id | human_gene_symbol | e-val | function | human reproductive mRNA expression? | human reproductive protein expression? | found on *Nautilus* Z? |
|---|---|---|---|---|---|---|---|
| obimac_0008617.1 | NP_001121055.1 | ING4 | 1.00E-109 | tumor suppressor protein that contains a PHD-finger | ovary, uterus, placenta, prostate, testis | ovary (fetal), testis, testis (fetal) | 0 |
| obimac_0008634.1 | NP_001182257.1 | RAB9A | 2.00E-92 | GDP binding activity; GTP binding activity; and GTPase activity. | ovary, uterus, placenta, prostate, testis | uterus, cervix, ovary, ovary (fetal), testis, testis (fetal) | 1 |
| obimac_0008637.2 | NP_001007469.1 | SMARCB1 | 0.00E+00 | SWI/SNF Related, Matrix Associated, Actin Dependent Regulator Of Chromatin, | ovary, uterus, placenta, prostate, testis | uterus, ovary, ovary (fetal), testis, testis (fetal) | 0 |
| obimac_0008638.1 | no hit | - | - | - | - | - | 0 |
| obimac_0008640.1 | NP_006310.1 | CDIPT | 8.00E-81 | Catalyzes the biosynthesis of phosphatidylinositol | ovary, uterus, placenta, prostate, testis | uterus, cervix, ovary, ovary (fetal), testis, testis (fetal) | 0 |
| obimac_0008642.1 | no hit | - | - | - | - | - | 1 |
| obimac_0008643.1 | NP_005207.3 | DDOST | 8.00E-128 | Subunit of the oligosaccharyl transferase (OST) complex | ovary, uterus, placenta, prostate, testis | uterus, cervix, ovary, ovary (fetal), testis, testis (fetal), seminal vesicle | 0 |
| obimac_0008647.1 | NP_003123.2 | SRM | 2.00E-114 | Spermidine Synthase | ovary, uterus, placenta, prostate, testis | uterus, cervix, ovary, ovary (fetal), testis, testis (fetal), seminal vesicle | 0 |
| obimac_0008648.1 | NP_001011547.2 | SLC5A9 | 0.00E+00 | Electrogenic Na(+)-coupled sugar symporter | ovary, uterus, prostate, testis | none | 1 |
| obimac_0008657.1 | NP_004152.1 | RAB1A | 9.00E-134 | Ras superfamily of GTPases. | ovary, uterus, placenta, prostate, testis | uterus, cervix, ovary, ovary (fetal), testis, testis (fetal), seminal vesicle | 1 |
| obimac_0008668.1 | NP_001287785.1 | FOSL1 | 2.00E-13 | FOS Like 1, AP-1 Transcription Factor Subunit | ovary, uterus, prostate, testis | ovary | 0 |
| obimac_0008926.1 | AAH14937.1 | FHIP1A | 3.00E-129 | protein localization to perinuclear region of cytoplasm | ovary, uterus, prostate, testis | testis (fetal) | 1 |
| obimac_0008950.6 | XP_005257828.1 | SPAG9 | 0.00E+00 | Sperm Associated Antigen; cancer testis antigen gene family | ovary, uterus, placenta, prostate, testis | uterus, cervix, ovary, ovary (fetal), testis, testis (fetal) | 1 |
| obimac_0008956.1 | no hit | - | - | - | - | - | 1 |
| obimac_0008962.2 | NP_002755.1 | PRPS1 | 0.00E+00 | catalyzes the phosphoribosylation of ribose 5-phosphate to 5-phosphoribosyl-1-pyrophosphate | ovary, uterus, placenta, prostate, testis | uterus, cervix, ovary, ovary (fetal), testis, testis (fetal) | 0 |
| obimac_0008963.1 | NP_005330.1 | UBE2K | 9.00E-89 | ubiquitin-conjugating enzyme | ovary, uterus, placenta, prostate, testis | uterus, cervix, ovary, ovary (fetal), testis, testis (fetal), seminal vesicle | 0 |
| obimac_0008964.1 | NP_001073308.1 | OCIAD1 | 9.00E-22 | OCIA Domain Containing 1; Maintains stem cell potency | ovary, uterus, placenta, prostate, testis | uterus, cervix, ovary, ovary (fetal), testis, testis (fetal) | 0 |
| obimac_0008989.8 | NP_001381782.1 | KLC1 | 0.00E+00 | Kinesin Light Chain 1 | ovary, uterus, placenta, prostate, testis | | 1 |
| obimac_0008993.1 | NP_001960.2 | EIF5 | 8.00E-147 | Eukaryotic translation initiation factor-5 | ovary, uterus, placenta, prostate, testis | uterus, cervix, ovary, ovary (fetal), testis, testis (fetal) | 1 |

Table S6: Table of blast results from orthologous region genes of chromosome Z to humans.

| **Species** | **Source** | **Accession ID** | **Citation** |
|---|---|---|---|
| *Nautilus pompilius* | CNCB | GWHBECW00000000 | Zhang et al. (2021) |
| *Nautilus pompilius* | www.doi.org/10.6084/ m9.figshare.14236208 | N/A | Huang et al. (2022) |
| *Euprymna scolopes* | NCBI | GCA_024364805.1 | Albertin et al. (2022a) |
| *Architeuthis dux* | NCBI | GCA_006491835.1 | Da Fonseca et al. (2020) |
| *Sepia pharaonis* | NCBI | GCA_903632075.3 | Song et al. (2021) |
| *Sepia esculenta* | Darwin Tree of Life | GCA_964036315.1 | N/A |
| *Octopus minor* | gigadb | N/A | Kim et al. (2018) |
| *Hapalochlaena maculosa* | NCBI | GCA_015501135.1 | Whitelaw et al. (2020) |
| *Octopus bimaculoides* | NCBI | PRJNA1076263 | This Article |
| *Octopus sinensis* | NCBI | GCF_006345805.1 | Li et al. (2020) |
| *Eldone cirrhosa* | Darwin Tree of Life Project | GCA_964016885.1 | N/A |

Table S7: Source of whole genome assemblies by species and the associated publications.
